# Evolutionary analysis of paired box gene family and biological function exploration of *Lr.Pax7* in lamprey*(Lethenteron reissneri)*

**DOI:** 10.1101/2024.05.06.592610

**Authors:** Ayqeqan Nurmamat, Zihao Yan, Yao Jiang, Haoran Guan, Ruyu Zhuang, Shuyuan Zhang, Yuesi Zhou, Min Xiu, Ya Pang, Ding Li, Liang Zhao, Xin Liu, Yinglun Han

**Author notes:** Correspondence (Y.H.), (X.L.). These authors contributed equally to this work (Ayqeqan Nurmamat, Zihao Yan, Yao Jiang).

## Abstract

The *Pax(paired box)*gene family comprises highly conserved transcription factors that play a crucial regulatory role in embryonic development, tissue morphogenesis, and cell differentiation. The gene family comprises of nine members, namely *Pax1* to *Pax9*, each exhibiting distinct functionalities and expression patterns. Among the members of the Pax family, *Pax7* is particularly involved in key regulatory functions related to skeletal muscle development and satellite cell biology in vertebrates. However, the precise role of *Pax7* and its downstream targets in jawless vertebrates such as lamprey remains relatively understudied. Therefore, investigating the expression and function of lamprey’s *Pax* genes holds significant value as a model organism for understanding developmental processes, evolution, and interactions with other vertebrates. In this study, we identified *Pax7* in *Lethentero reissneri* and multiple bioinformatics analyses suggested that it may represent the ancestral gene of the *Pax7* shared among higher vertebrates. To further elucidate the function and regulatory mechanism of *Pax7*, we performed transcriptome sequencing analysis after silencing *Lr.Pax7* expression which revealed alterations in metabolic processes along with associated genes(*Plcy,Jagged2,Hes,Myod,Frizzled2*,etc) involved in various immune pathways(*Notch, Wnt, IL-17, TNF*, etc), myogenesis process, adipocyte formation process consistent with those observed in higher vertebrates. Comparative studies between lamprey and high vertebrates could provide valuable insight into understanding the functional role of *Pax7* and its downstream targets across diverse biological process as well as disease states.

## Introduction

Gene regulation refers to the precise regulation of gene expression in an organism, and transcription factors are proteins that bind to DNA and regulate gene expression by promoting or inhibiting the expression of target genes. From the late 1980s[1], scientists started studying special genes called Pax genes that control how genes work in organisms as they grow. There are nine Pax genes found in animals like mice, zebrafish, and humans[2]. Based on the composition domain and homology the sequence the *Pax* family divided into four subfamilies *Pax1/9, Pax2/5/8, Pax4/6, Pax3/7[3]. Pax7*plays a pivotal role in the implementation, protection, and repair of skeletal muscle. *Pax7* helps to control the balance between self-renewal and differentiation of satellite cells, ensuring that they can proliferate when needed to generate new muscle cells and differentiate into mature muscle fibers when necessary for muscle development and repair. The expression of the *Pax7* gene in nerve cells is critical for dorsal root and sensory ganglia development. The *Pax7* gene serves as a primary controlling factor for skeletal muscle development while influencing different biological processes; however, its exact role remains limited regarding jawless vertebrates like lamprey requiring extensive research to comprehend intricate mechanisms involved. Given the lamprey’s unique status as an ancient jawless fish, possessing an ancient lineage and distinctive biological features, it presents a rare opportunity to explore gene function across hundreds of millions of years of vertebrate evolution. Utilizing the lamprey as a model system for gene function research represents an innovative approach in the fields of evolutionary and comparative genomics. In this study, we investigated the regulatory mechanism of *Pax7* in lamprey by gene cloning, gene expression analysis, gene silencing and transcriptome data analysis. We also explored the interaction between genes with significant differences. Identification of *Lr.Pax7* in lamprey tissues began with retrieving protein sequences that are similar to human Pax family members in sea lamprey (*Petromyzon marinus*) or zebrafish (*Danio rerio*) from the NCBI protein database and employing the BLAST to identify corresponding homologs. Subsequently, we extracted the Pax sequences from our library. *Lethenteron reissneri* specimens were dissected to isolate various tissues. Designed primers based on *Pax7* nucleotide sequences in lampreys’ cDNA library targeting pax domains, aiming to verify the effectiveness of lampreys’ cDNA as a template for validation (Supplementary Table S3). *Lr.Pax7* was successfully amplified via PCR in muscle tissue.

## Methods and results

Here a variety of methods are used to analyze bioinformatics, The results show that the amino acid sequence of *Pax7* shows high similarity among animals (Figure 1A), with a decreasing trend from higher to lower organisms, as revealed by sequences alignment. It can be observed from the evolutionary tree (Figure 1B) that pax genes for each subfamily are present in ancestral chordate and that pax genes are present in amphioxus. The *Petromyzon marinus, Lethenteron camtschaticum, Lethenteron reissneri* constitute a sister group, which has become a good model for the study of jawless vertebrates.*Pax9,Pax2,Pax6*, and *Pax7* show high similarity to those of other higher vertebrates. Therefore, these genes were named as *Lr.Pax9, Lr.Pax2, Lr.Pax7*, and *Lr.Pax6*. The results indicate that the *Pax7* gene is significantly preserved across various species, from higher to lower. This suggests that the DNA sequence of the gene is remarkably similar among different species. *Lr.Pax7* is positioned between vertebrates and invertebrates and is most closely related to *P. marinus Pax7*. To further investigate the evolutionary history of *Pax7* in vertebrates, we compared the genetic environment of *Pax7* (Figure 1C). This gives more insight into the original evolutionary position of the lamprey. The crystal structure prediction analysis shows that *Lr.Paxs* and *Hm.Paxs* have highly homologous structure (Figure 1D). The *Pax* gene has a similar structure (Figure 1E), including a conserved DNA-binding structure called pair-box domain. This structure contains about 128 amino acids and is responsible for binding specific DNA sequences, regulating gene expression, and interacting with other proteins. In addition, many *Pax* gene members also contain DNA-binding structures called homeodomains, which play an important role in gene expression regulation and the development and establishment of various tissues and organs. *Pax* genes may also have domains that regulate transcriptional activation or inhibition, interacting with other proteins to enhance or inhibit transcription of target genes. Some motifs were detected in all sequences, indicating that they are the most primitive and conserved functional motifs. This phenomenon also suggests that these motifs may perform similar biological functions in different species, even though they may have varied to some extent in different branches of evolution. Thus, the conservation of such motifs (Figure 1F) may mean that they are essential for specific biological functions that may be required across different species, leading to the retention of these motifs. Overall, the structure of *Pax* genes includes elements such as pairing box domains and homologous domains, which enable *Pax* genes to bind specific DNA sequences and play important roles in embryonic development and tissue differentiation. The pax gene family domain is also shown in the figure. It can be seen that the domain of *Lr.Pax7* is basically consistent with that of higher vertebrates, which further proves the conserved nature of *Pax7* gene. The gene *Lr.Pax7* serves as the ancestral form of *Pax7* in vertebrates, and our study carries significant evolutionary implications.

**Figure 1.**
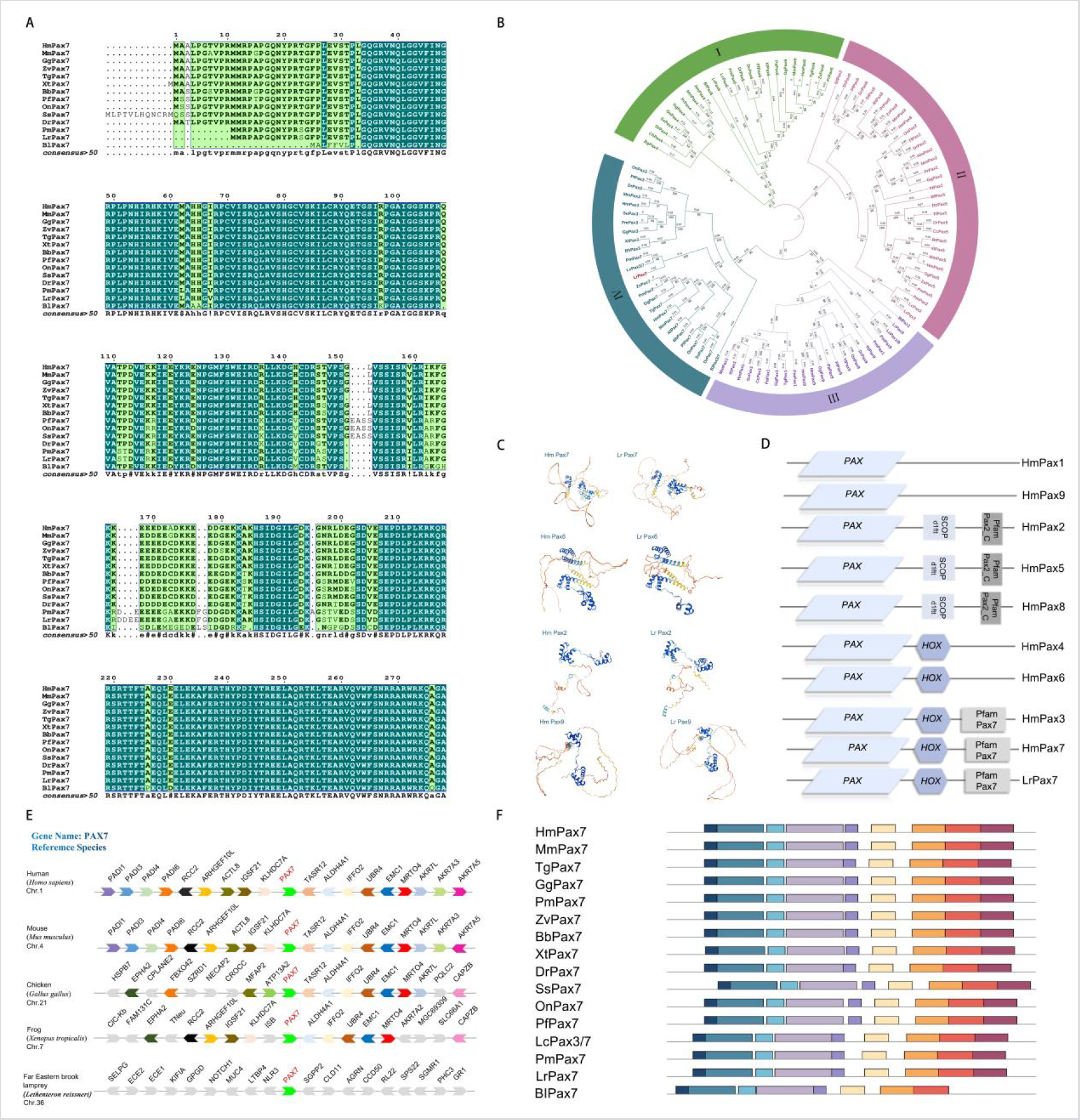
Bioinformatics analysis. The identification and sequence analysis of Pax family genes. (A) Protein sequences alignment from *Hm.Pax7-Bl.Pax7*. (B) The phylogenetic tree for the Pax family based on the NJ method. (C) Diverse genomic synteny analysis from *Hm.Pax7* to *Lr.Pax7*. (D)Prediction of the tertiary structure of *Pax7,Pax6,Pax2,Pax9* in human and lampreys. (E) comparison of the domains between *Pax1-Pax9* from humans and lampreys. (F) A motif composition of the *Pax7* family proteins. The full motif composition of Pax family and the full names of species are listed in Supplementary Table 1, the matching amino acid sequences in motifs are listed in Supplementary Table 2.

We also used both qPCR and Western Blotting (WB) methods to detect the expression of a gene in various tissues to study the expression pattern and level of the gene in different tissues. The expression patterns of *Pax7* in adult *Lethenteron reissneri* were investigated by conducting qPCR analysis on total RNA samples extracted from five normal tissues, namely gill, kidney, marrow, brain, and muscle. In western blotting, the band density is significantly higher in the muscle sample compared to the other tissues, indicating robust expression in the muscle tissue (Figure 2A). The results revealed that *Lr.Pax7* exhibited predominant expression in muscle tissue, followed by kidney, gill, marrow and brain (Figure 2B). In order to better understand the biological function of *Lr.Pax7*, specifically target the pax domain of *Lr.Pax7*, the siRNA sequences were carefully selected to ensure efficient and specific knockdown of the *Lr.Pax7* gene. Following synthesis, the siRNA was administered to lamprey specimens as part of an experimental study (Supplementary TableS4). Ten lampreys, each weighing approximately 4 g, were selected for the study. The lampreys were divided into two groups: the experimental group (SiRNA) and the negative control group (NC). The results demonstrated a particularly effective silencing efficiency of muscle tissue at 72 hours post-treatment (Figure 2C). Previous studies have indicated the presence of *Pax3* and *Pax7* genes in adult muscle tissues of higher vertebrates. These observations suggest tha*t Pax* derived genes may retain ancestral functions. Subsequently, we selected muscle tissue for transcriptome analysis.The tissues exhibiting superior silencing efficiency (three biological replicates) were submitted to oebiotech for transcriptome sequencing. We mapped the volcano based on the differential genes (Supplementary Figure1A), a total of 630 genes were up-regulated while 839 genes were down-regulated (Supplementary Figure1B). Then, conducted enrichment analysis on these differentially expressed genes, and the results are as follows. 10 bases with high significance and related to cancer pathways and *PPAR* pathway were selected (Supplementary TableS5). Subsequently, qPCR experiments were performed with GAPDH as the reference gene to verify the authenticity of the transcriptome results. Experimental results All homogeneous transcriptome results were consistent (Figure 2D), proving the validity of the transcriptome data. According to the functional classification, the KEGG is usually divided into three levels. The GO top 20 results showed that the changes of ribosomes and collagen in the cell components were obvious during gene interference. Significant changes were observed in oxidation-reduction, metabolic process and translation. The activity of oxidoreductase, structural consistent of ribosome, the composition of extracellular structure matrix and the catalytic activity in the functional part of the molecule changed significantly (Figure 2E-Figure 2G). KEGG top 20 enrichment analysis using bubble maps revealed significant enrichment of differential genes in *Th1* and *Th2* differentiation, *HIF-1*, and *IL-17*. Analysis of KEGG 1-3 levels revealed significant changes involved in the *Wnt,Notch*, and *PPAR* signaling pathways during *Pax7* silencing in lampreys. Some of the up-regulated genes have been linked to the activation of oncogenes or dysregulation of signaling pathways that lead to the onset and progression of cancer (Figure 2H-Figure 2J).

**Figure 2.**
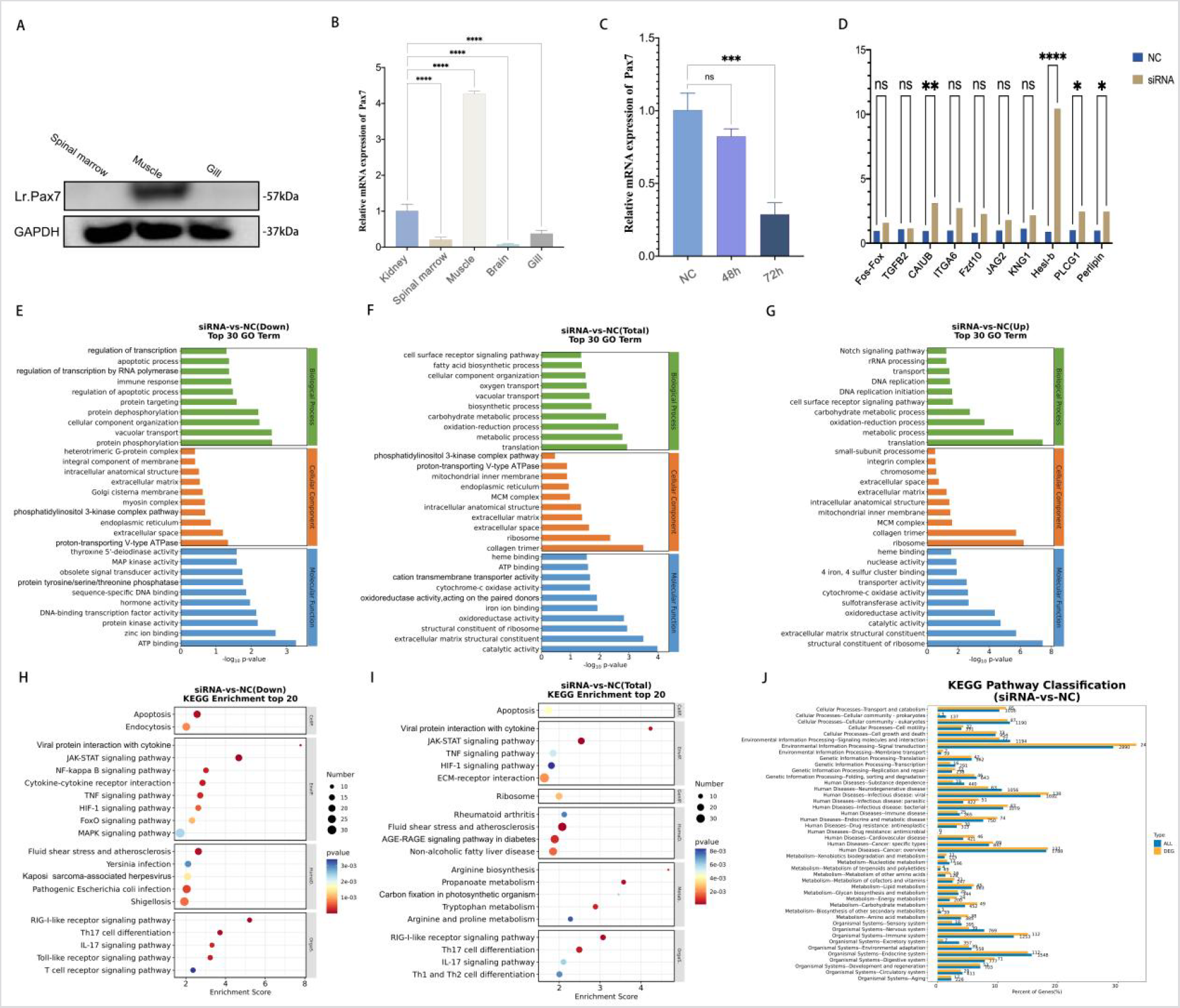
Expression of *Lr.Pax7* in tissues and analysis of *Lr.Pax7* Gene silencing Transcriptome data. (A) Relative *Lr.Pax7* mRNA expression levels in various lamprey tissues. (B) Western blot analysis of protein expression levels of *Lr.Pax7* in different tissues. (C) The expression of *Lr.Pax7* gene was quantified using qPCR following silencing of the *Lr.Pax7* gene, and an abundance map was generated. *Lr.GAPDH* served as the reference gene, and the experimental results were expressed as means ± SD. ^*^Indicates significant difference compared with the control group (n = 3, Ns: not significant, ^*^ P < 0.05, ^**^ P < 0.01,^***^ P < 0.005. (D) Bar chart illustrating the detection of relative expression levels of differential genes in the *Lr. Pax7* transcriptome using Q-PCR. Ns denotes non-significant, ^*^P < 0.05, ^**^P < 0.01, ^***^ P < 0.005 and ^****^ P < 0.001 (paired t-test). (E) Go enrichment analysis results of *Lr.Pax7* Gene silencing Transcriptome Down-regulated expression genes. (F) Go enrichment analysis results of *Lr.Pax7* Gene silencing Transcriptome Total-regulated expression genes. (G) Go enrichment analysis results of *Lr.Pax7* Gene silencing Transcriptome Up-regulated expression genes. (H) KEGG enrichment analysis results of *Lr.Pax7* Gene silencing Transcriptome Down-regulated expression genes.(I) KEGG enrichment analysis results of *Lr.Pax7* Gene silencing Transcriptome Total-regulated expression genes. (J) KEGG pathway classification results of *Lr.Pax7* Gene silencing Transcriptome.

The silencing of *Lr.Pax7* in lampreys led to a series of changes affecting metabolism and immune processes, as evidenced by Go and KEGG pathway analyses. Suppression of *Lr.Pax7* influenced key metabolic enzymes, suggesting a role in vital metabolic pathways and triggering a stress response impacting energy metabolism, hormone levels, and redox balance. Signaling molecules, such as *NFKBIA, TGFBR1, Fos, Bcl2, Bcl9*, and *Bcl10*, were activated, implicating *Lr.Pax7* in the lamprey’s immune response and inflammation. Upregulated genes, especially *NFKBIA* and *TGFBR1*, point towards modulation of *NF-κB* and *TGF-β* pathways. Moreover, the upregulation of Fos further underscores the influence on immune response genes. After observing the differentially expressed genes based on the function of *Pax7* in higher vertebrates, it was found that the *Pax7* related genes (*Myog, Six4, Six1 and MyoD*) were also changed in lamprey and could be found in the data. *Pax7, Myog, Six4, Six1* and *MyoD* play pivotal roles in muscle growth and differentiation[4-6]. *Pax7* serves as a crucial regulator of muscle stem cells (satellite cells) and is indispensable for maintaining an undifferentiated myogenic precursor cell pool. Their collective actions form a regulatory network essential for muscle cell formation. This suggests not only that *Lr.Pax7* may be involved in the regulation of muscle development and differentiation in lamprey, but also that the function of *Pax7* in higher vertebrates may be conserved in this ancient jawless fish. Interestingly, silencing *Lr.Pax7* leading to lower *PDGFRα,PPARγ,Fabp4* expression and hindered lipid droplets. These findings collectively highlight *Lr.Pax7*’s significant regulatory role in lamprey immunity, inflammation, and muscle development, paralleling its function in advanced vertebrates.

## Discussion

Taken together, in this study a combination of diverse bioinformatics analyses, experimental investigations, transcriptome analyses, and other methodologies were employed to further elucidate the evolution and biological functionality of *Pax7*. The phylogenetic tree demonstrated that the *Pax* family was classified into four distinct subfamilies, consistent with previous studies. *Lr.Pax7is* positioned in the outgroup of vertebrates within the phylogenetic tree. This observation suggests an evolutionary relationship from primitive to advanced forms in lampreys and implies that *Lr.Pax7* genes may represent ancestral *Pax7* gene in vertebrates. Furthermore, comprehensive domain similarity analysis indicates that *Lr.Pax7* likely shares similar functions with its counterparts in other vertebrate species, highlighting a high degree of conservation across different organisms. Additionally, this study delves into exploring potential conserved regulatory mechanisms underlying the evolution of *Pax7*. The conservation of molecular pathways between lamprey and human suggests that fundamental genetic regulatory mechanisms were established early in vertebrate evolution and have remained conserved throughout the evolutionary process. This similarity provides valuable insights into the molecular and cellular processes involved in stem cell regulation, tissue regeneration, as well as underlying pathways associated with diseases. These findings suggest that *Pax7* plays a critical regulatory role in various biological processes in lampreys, including immune-related processes and metabolism. The dysregulation of *Pax7* may contribute to the development and progression of various diseases, including cancer. Further studies are needed to fully elucidate the molecular mechanisms underlying the role of *Pax7* in these processes.

## Author Contributions

Conceptualization: Y.H. and X.L.; Methodology: Y.H. and X.L.; Investigation: Y.Z., Z.Y., G.H.,M.X. and Y.Z.; Funding acquisition: Y.H. and X.L; Project administration: S.Z., R.Z., D.L., L.Z. and Y.P.; Writing—original draft: Y.Z., A.N. ; Writing—review and editing: A.N., Z.Y., Y.H. and X.L. All authors have read and agreed to the published version of the manuscript.

