## Supplementary material for "Evolutionary analysis of paired box gene family and biological function exploration of *Lr.Pax7* in lamprey*(Lethenteron reissneri)*": 11Supplementary_Materials-2.pdf

### Supplementary Materials

#### Method and materials

##### Experimental animals

The lampreys utilized in this study were *Lethenteron reissneri* specimens weighing 3-5 g, with approximately 12 individuals employed for the experimental procedures. These lampreys were bred in the Tongjiang Valley of the Songhua River in Heilongjiang Province, China, and subsequently reared in a culture channel with circulating water at a constant temperature of 4°C. The handling of lampreys and all experimental protocols were approved by the Ethics Committee for Animal Welfare and Research of Liaoning Normal University and Dalian Medical University (Permit number: SYXK2004−0029).

Identification of *Pax* family and cloning of *Lr.Pax7*

We retrieved protein sequences homologous to human members of the Pax family in sea lamprey (*Petromyzon marinus*) or zebrafish (*Danio rerio*) from the NCBI protein database (https://www.ncbi.nlm.nih.gov). These sequences were utilized as queries for a BLAST search against our personal lamprey database, employing the NCBI's Basic Local Alignment Search Tool (BLAST), to identify corresponding homologs. Subsequently, we extracted the Pax sequences from our library. *Lethenteron reissneri* specimens were dissected to isolate tissues such as heart, liver, intestine, kidneys, gills, muscles, spinal cord and others. mRNA was then extracted and reverse-transcribed into cDNA. Using the *Pax7* nucleotide sequence present in the Lampreys cDNA library as a template, primers containing *pax* domain were designed to validate the efficacy of lampreys cDNA antitemplate. *Lr.Pax7* was successfully amplified via PCR in muscle tissue.

**Bioinformatics analysis of the *Pax* family**

Phylogeny analysis, conserved motifs and domains analysis

The protein sequences of the *Pax* family were carefully compared using BioEdit sequence alignment editor. Subsequently, MEGA7.0 software was employed to construct a phylogenetic tree to elucidate the evolutionary relationships within the *Pax* family. To ensure statistical robustness during the tree construction process, 1000 bootstrap replicates were performed using the Neighbor-joining method. Conserved motifs in Pax proteins were identified and analyzed using MEME 5.1.1, while SMART tools were utilized for online systematic analysis of conserved domains in Pax proteins. Our research findings contribute to understanding the inherent conserved structural elements within the Pax protein family, which are visually presented through ITOL (Interactive Tree of Life, https://itol.embl.de/itol.cgi) platform, providing valuable insights into their evolutionary history and relevance, as well as facilitating a deeper understanding of their evolutionary conservation and interspecies variation.

Synteny construction, crystal structure prediction, prediction of protein sequence alignment

The genomic collinearity was established using the Genomics online tool (https://www.genomicus.bio.ens.psl.eu/genomicus-110.01/cgi-bin/search.pl). Analysis encompassed multiple species, including human, rat, chicken, frogs, and lampreys. Crystal structures were predicted using the SWISS-MODEL software, in accordance with established protocols for computational modeling of protein structures. To align the *Pax7* protein sequences across various species, we employed the online ESPript3.0 online tool (https://espript.ibcp.fr/ESPript/ESPript/).

#### qRT-PCR and Western Blotting Analysis of *Lr.Pax7*

First, total RNA was extracted from the heart, liver, intestine, kidney, brain, gill, and muscle tissues of lampreys. Subsequently, the extracted RNA was utilized as a template for cDNA synthesis using reverse transcription kits to convert mRNA into cDNA. The expression of *Lr.Pax7* in each tissue was then analyzed using a qPCR kit (RR820A, TaKaRa), with the cDNA from each tissue serving as a template. Specific primers for Lr-Pax7 were meticulously designed using PrimerPremier5.0 software. The expression of Pax mRNA was determined by employing the synthesized cDNA. Quantitative reverse transcription PCR (qRT-PCR) was performed on a PCR Thermal Cycler Dice Real-Time System (TaKaRa, Japan). Each reaction had a volume of 20 μL and included 2 μL cDNA (50 ng/μL), 0.4μL 10 Mm reverse primer, 0.4 μL 10 Mm forward primer, 10 μL 2 × SYBR premix EX Taq (TaKaRa, Japan), and 7.2 μL dd$H_2O$.The real-time fluorescent PCR reaction followed specific conditions including initial denaturation at 95℃ for 30 s, during which denaturation occurred at 95℃ for 5 s, followed by annealing at 55℃ for 30 s, and extension at 72℃ for 30 s. This process underwent 40 cycles. All reactions were repeated three times to ensure accuracy and reliability while utilizing *Lr-GAPDH* as an internal control. In western blotting, the protein extracted from muscle, gill, and marrow tissue was mixed with loading buffer, denatured at 100°C for 5 minutes using the BCA Protein Assay kit (Cat). No. 5000001; BioRad, Hercules, CA, USA), subjected to SDS-PAGE based on target protein's relative molecular mass and transferred onto nitrocellulose membranes. After blocking with 5% nonfat dry milk in PBS for 2 hours at room temperature, the membranes were incubated overnight at 4°C with primary antibodies against Pax7 and GAPDH (Sunon Biotechnology, China), respectively. Following washing with diluted phosphate buffer solution (PBS) (0.1 %), cells were incubated for an hour at room temperature with HRP-labeled goat anti-rabbit IgG(1:500). Finally, Western blot bands were quantified using an Odyssey Infrared Imaging system (LI-COR) and Odyssey v3.0 software.

#### Gene silencing and experimental validation

To specifically target the pax domain of *Lr.Pax7*, small interfering RNA (siRNA) was designed and synthesized by Genepharma company. The siRNA sequences were carefully selected to ensure efficient and specific knockdown of the *Lr.Pax7* gene. Following synthesis, the siRNA was administered to lamprey specimens as part of an experimental study. Ten lampreys, each weighing approximately 4 g, were selected for the study. The lampreys were divided into two groups: the experimental group (SiRNA) and the negative control group (NC). Intraperitoneal injection was performed, with a dose of 30 μl/g and a concentration of 53.67 μg/mol for both groups. Subsequently, tissue samples were collected at two specific time intervals (48 hours and 72 hours) post-injection and immediately frozen to preserve the RNA content. Total RNA was then extracted from the collected tissues, and reverse transcription was carried out using appropriate reagents and reverse transcriptase to convert the RNA into complementary DNA (cDNA).For quantitative analysis of gene expression, the cDNA obtained from the muscle tissue was used as the template for the quantitative real-time PCR (qPCR) reaction. The target gene, *Lr.Pax7*, and the internal reference gene, *Lr.GAPDH*, were amplified with three replicates each in the muscle tissue samples. The expression level of the internal reference gene, *Lr.GAPDH*, served as the baseline for normalization, and the data from the control group (NC) were used as the reference for comparison. The relative expression level of the *Lr.Pax7* gene was calculated using the *2-ΔΔCT* method, allowing for accurate quantification of gene expression changes in response to siRNA treatment.

#### Transcriptome sequencing

After the tissue with the successfully suppressed genes was submitted to oebiotech(Shanghai, China) company for transcriptome sequencing, the obtained results were thoroughly analyzed. Specifically, 10 differential genes were identified from both the KEGG (Kyoto Encyclopedia of Genes and Genomes) and GO (Gene Ontology) databases. These genes were selected based on their potential significance in the context of the study and were then subjected to primer design for further analysis. The designed primers were tailored to specifically amplify each of the selected genes. Following this, the muscle tissue collected at the 72-hour time point post-suppression was utilized as a template for qPCR (quantitative polymerase chain reaction) verification.

#### Statistical analysis

The bar chart data in this study were subjected to analysis using three independent replicates of existing data. We calculated the standard deviation and established confidence intervals (mean ± standard deviation (SD)), along with conducting statistical tests (two-tailed t-test). Due to the significant stress response of larval lampreys to siRNA or stimuli injected intraperitoneally, an initial sample size of 10 was chosen. However, a minimum of three samples was maintained, and if the sample size dropped below three, the experiment was repeated. All statistical analyses were performed using GraphPad Prism 8.0 software. Differences between treatment groups were assessed through a two-way analysis of variance (ANOVA). A significance threshold of $P < 0.05$ ($^{*}P < 0.05$, $^{**}P < 0.01$, $^{***}P < 0.005$, $^{****}P < 0.001$) was set and adjusted for multiple comparisons using the Bonferroni correction method.

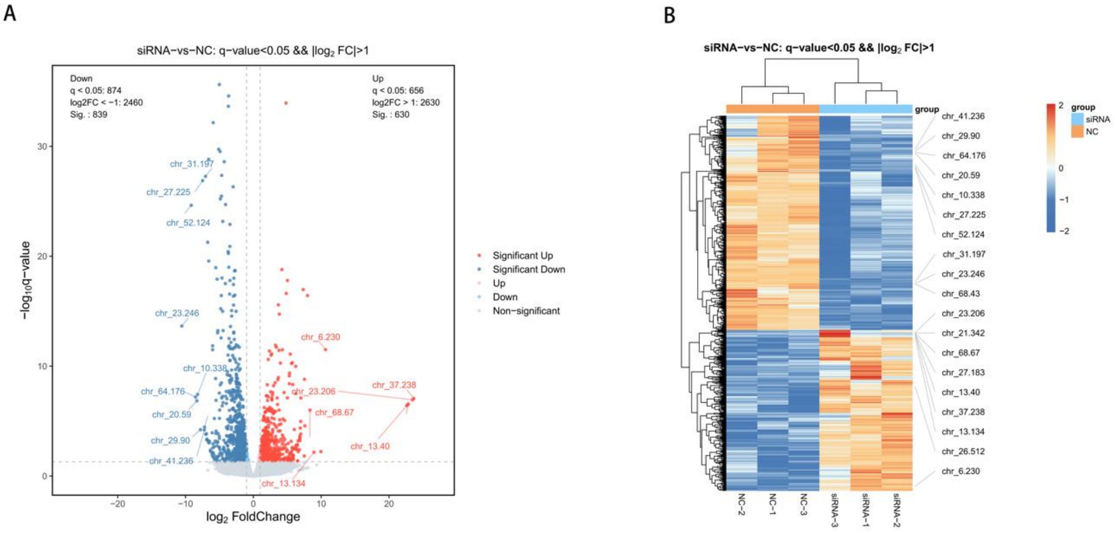


**Supplementary Figure1. Enrichment analysis** （A) A volcano map drawn from differential gene analysis .（B）Up-down-regulated gene analysis diagram between NC and SiRNA groups.

#### **Table S1** Search for the full species name of the Pax gene

| Symbol | Species | Symbol | Species | Symbol | Species |
|---|---|---|---|---|---|
| HmPax9 | Homo sapiens | OsPax6 | Octopus sinensis | StPax8 | Salmo trutta |
| MmPax9 | Mus musculus | HmPax3 | Homo sapiens | HmPax2 | Homo sapiens |
| GgPax9 | Gallus gallus | MmPax3 | Mus musculus | MmPax2 | Mus musculus |
| XtPax9 | Xenopus tropicalis | GgPax3 | Gallus gallus | GgPax2 | Gallus gallus |
| PaPax9 | Protopterus annectens | PmPax3 | Petromyzon marinus | PrPax2 | Podarcis raffonei |
| RtPax9 | Rana temporaria | SuPax3 | Sceloporus undulatus | OnPax2 | Oreochromis niloticus |
| DrPax9 | Danio rerio | BbPax3 | Bombina bombina | TfPax2 | Tachysurus fulvidraco |
| OnPax9 | Oreochromis niloticus | XtPax3 | Xenopus tropicalis | DrPax2 | Danio rerio |
| TfPax9 | Tachysurus fulvidraco | PrPax3 | Podarcis raffonei | XtPax2 | Xenopus tropicalis |
| PmPax9 | Petromyzon marinus | PfPax3 | Perca flavescens | PmPax2 | Petromyzon marinus |
| HmPax1 | Homo sapiens | OnPax3 | Oreochromis niloticus | Lc2/5/8 | Lethenteron camtschaticum |
| MmPax1 | Mus musculus | LcPax3/7 | Lethenteron camtschaticum | LrPax2 | Lethenteron reissneri |
| XtPax1 | Xenopus tropicalis | BlPax3/7 | Branchiostoma lanceolatum | BfPax2 | Branchiostoma floridae |
| PaPax1 | Protopterus annectens | HmPax7 | Homo sapiens | | |
| CcPax1 | Cyprinus carpio | MmPax7 | Mus musculus | | |
| SsPax1 | Salmo salar | TgPax7 | Taeniopygia guttata | | |
| GgPax1 | Gallus gallus | GgPax7 | Gallus gallus | | |
| TgPax1 | Taeniopygia guttata | PmPax7 | Petromyzon marinus | | |
| PrPax1 | Podarcis raffonei | ZvPax7 | Zootoca vivipara | | |
| LcPax1/9 | Lethenteron camtschaticum | BbPax7 | Bombina bombina | | |
| LrPax1 | Lethenteron reissneri | XtPax7 | Xenopus tropicalis | | |
| PmPax1 | Petromyzon marinus | DrPax7 | Danio rerio | | |
| BfPax1 | Branchiostoma floridae | SsPax7 | Salmo salar | | |
| HmPax4 | Homo sapiens | OnPax7 | Oreochromis niloticus | | |
| MmPax4 | Mus musculus | PfPax7 | Perca flavescens | | |
| PrPax4 | Podarcis raffonei | PmPax7 | Petromyzon marinus | | |
| GgPax4 | Gallus gallus | LrPax7 | Lethenteron reissneri | | |
| DrPax4 | Danio rerio | HmPax5 | Homo sapiens | | |
| ChPax4 | Clupea harengus | MmPax5 | Mus musculus | | |
| SsPax4 | Salmo salar | GgPax5 | Gallus gallus | | |
| OnPax4 | Oreochromis niloticus | ZvPax5 | Zootoca vivipara | | |
| BgPax4 | Bufo gargarizans | PmPax5 | Petromyzon marinus | | |
| HmPax6 | Homo sapiens | RtPax5 | Rana temporaria | | |
| MmPax6 | Mus musculus | XlPax5 | Xenopus laevis | | |
| ZvPax6 | Zootoca vivipara | TfPax5 | Tachysurus fulvidraco | | |
| PrPax6 | Podarcis raffonei | DcPax5 | Denticeps clupeoides | | |
| GgPax6 | Gallus gallus | DrPax5 | Danio rerio | | |
| TgPax6 | Taeniopygia guttata | CcPax5 | Cyprinus carpio | | |
| DcPax6 | Zootoca vivipara | LcPax2/5/8 | Lethenteron camtschaticum | | |
| PfPax6 | Perca flavescens | HmPax8 | Homo sapiens | | |
| DrPax6 | Danio rerio | MmPax8 | Mus musculus | | |
| PaPax6 | Protopterus annectens | PrPax8 | Podarcis raffonei | | |
| XlPax6 | Xenopus laevis | ZvPax8 | Zootoca vivipara | | |
| LcPax6 | Lethenteron camtschaticum | RtPax8 | Rana temporaria | | |
| PmPax6 | Petromyzon marinus | DrPax8 | Danio rerio | | |
| LrPax6 | Lethenteron reissneri | PfPax8 | Perca flavescens | | |
| BfPax6 | Branchiostoma floridae | DcPax8 | Denticeps clupeoides | | |

**Table S2** The conserved motifs discovered among the amino acid sequences of the *Pax7* gene families from vertebrates and invertebrates using the MEME system.

| Number | Best possible match |
|---|---|
| 1 | QQRSRTTFTQEQLEALEKAFERTHYPDIY |
| 2 | FVNGRPLPBVIRQRIVELAHQGIRPCDISRQLRVSHGCVSK |
| 3 | RYYETGSIRPGAIGG |
| 4 | KPRVATPKVVKKIAEYKRZNPGMFAWEIRDRLLAEGVCDNDNVPSVSSIN |
| 5 | RILRNKVGQKS |
| 6 | SWPSSHSISGILGIRSSSEDE |
| 7 | GHGGVNQLGGV |
| 8 | TREELAQRTKLTEARVQVWFSNRRARWRKZA |
| 9 | GANQLAAFNHLJPGGFPPTGMPTLPPYQL |

**Table S3** *Lr-Pax7* PCR/Q-PCR Primer

| Gene name | Gene ID | PCR/RT- PCR | F-Primer | R-Primer |
|---|---|---|---|---|
| Lr-Pax7 | chr_36.238 | PCR | GACCTGTGGGTTCTCTAC | ACCAGGACAGAACTACCC |
| | | RT-PCR | GCTGCGTTACGTTCTGCAT | GTGTGCTCTTCTCTGCCGAT |

**Table S4** siRNA sequences

| Gene name | Sense（5'-3'） | Antisense（5'-3'） |
|---|---|---|
| Pax7-1 | GGGUAAAGAAGCUGCGUUATT | UAACGCAGCUUCUUUACCCTT |
| Pax7-2 | GAAUGUGGAAUUUCCACCATT | UGGUGGAAAUUCCACAUUCTT |
| Pax7-3 | CAGGAAUGGACCAAUUAAATT | UUUAAUUCGUCCAUUCCUGTT |
| Negative Control | UUCUCCGAACGUGUCACGUTT | ACGUGACACGUUCGGAGAATT |

**Table S5** Q-PCR Primer

| Gene name | F-Prime | |
|---|---|---|
| Fos-Fox | AGCAGCAGCACCACCATCAAC | GCAGCGGCTTGGCAGACTC |
| TGFB2 | GACGACGACGACGAGGAGGAG | AGACGGCGACGGGTTGGAG |
| CAIUB | CACAACACCACCTACGGCTTCC | ATCAACACCACCACCAACATCACC |
| ITGA6 | CCACTCTGGCAAGGAGAACTACAC | GGAGGAGGAGAAATGAGCGTGAAC |
| Fzd10 | GCTCCCTCGGTGGCTCCTG | GGTTGTTGTTGCTGCTGCTCTTG |
| JAG2 | CGACCACAACCACCACGATCAC | CTCCTCATCCTCGTCCTCCTCATC |
| KNG1 | GCTGTCCGAGTGTCTCTTCAAGTC | GCGTCTCTTCGAGGATTGGCTTC |
| Hesl-b | GCTGTCCGAGTGTCTCTTCAAGTC | GCGTCTCTTCGAGGATTGGCTTC |
| PLCG | GCCGTGTCCAAGTGCGTAGTG | GCTGCTTCTTGCCACCGTAGTC |
| Perlipin | GCCGTGTCCAAGTGCGTAGTG | GCTGCTTCTTGCCACCGTAGTC |
